# Deep spectral learning for label-free optical imaging oximetry with uncertainty quantification

**DOI:** 10.1101/650259

**Authors:** Rongrong Liu, Shiyi Cheng, Lei Tian, Ji Yi

**Affiliations:** Department of Biomedical Engineering, Northwestern University, Evanston, IL, 60208, USA; Department of Electrical and Computer Engineering, Boston University, Boston, MA 02215, USA; Department of Medicine, Boston University School of Medicine, Boston Medical Center, Boston MA, 02118, USA; Department of Biomedical Engineering, Boston University, Boston MA, 02215, USA

## Abstract

Measurement of blood oxygen saturation (sO_2_) by optical imaging oximetry provides invaluable insight into local tissue functions and metabolism. Despite different embodiments and modalities, all label-free optical imaging oximetry utilize the same principle of sO_2_-dependent spectral contrast from hemoglobin. Traditional approaches for quantifying sO_2_ often rely on analytical models that are fitted by the spectral measurements. These approaches in practice suffer from uncertainties due to biological variability, tissue geometry, light scattering, systemic spectral bias, and variations in experimental conditions. Here, we propose a new data-driven approach, termed deep spectral learning (DSL) for oximetry to be highly robust to experimental variations, and more importantly to provide uncertainty quantification for each sO_2_ prediction. To demonstrate the robustness and generalizability of DSL, we analyze data from two visible light optical coherence tomography (vis-OCT) setups across two separate *in vivo* experiments in rat retina. Predictions made by DSL are highly adaptive to experimental variabilities as well as the depth-dependent backscattering spectra. Two neural-network-based models are tested and compared with the traditional least-squares fitting (LSF) method. The DSL-predicted sO_2_ shows significantly lower mean-square errors than the LSF. For the first time, we have demonstrated *en face* maps of retinal oximetry along with pixel-wise confidence assessment. Our DSL overcomes several limitations in the traditional approaches and provides a more flexible, robust, and reliable deep learning approach for *in vivo* non-invasive label-free optical oximetry.

## 1. Introduction

Microvascular systems deliver oxygen to support cellular metabolism and maintain biological functions. Within the local microenvironment of blood vessels, oxygen unloads from hemoglobin and diffuses freely from red blood cells (RBC) to tissues following the gradient of oxygen partial pressure (pO_2_), which determines the oxygen saturation (sO_2_) of hemoglobin. The measurement of microvascular sO_2_ thus can provide invaluable insight into local tissue metabolism, inflammation, and oxygen-related pathologies (e.g. cancers, diabetic milieu and complications, and cardiovascular diseases) [1–5].

In recent years, several non-invasive and label-free optical imaging oximetry techniques have been developed to measure microvascular sO_2_. Despite their differences, the fundamental mechanism is the same that is based on the sO_2_-dependent spectral contrast from hemoglobin [6]. The spectral measurement is then related to sO_2_through a complex physical model incorporating tissue geometry, heterogeneous tissue scattering, light attenuation and propagation, and imaging optical instruments. This model is often simplified and analytically formulated under different approximations and assumptions. Examples include spatial frequency domain imaging [7,8] in the diffusive regime under the P3 approximation, multi-wavelength imaging [9–12] and visible light optical coherence tomography (vis-OCT) [13–26] in the ballistic regime based on the Beer’s law combined with the first Born approximation, photoacoustic microscopy/tomography assuming a uniform laser fluence inside the tissue [27,28], and photothermal imaging assuming a linear relation between the blood absorption and the change of the optical signal [29–32]. The sO_2_ estimation thus requires solving an ill-posed inverse problem that is inevitably subject to model inaccuracies, noise, systemic spectral bias, and experimental conditions. One widely used inversion method is the spectral least-squares fitting (LSF) that estimates the sO_2_ by matching the spectral data with the analytical model. However, inpractice multiple sources of spectral errors exist that are impossible to be fully parameterized in a simple analytical form, which in turn compromises the sO_2_ estimation accuracy, repeatability, and cross-comparison between different devices, test subjects, and time. Therefore, it is imperative to develop a more robust model to enable more accurate quantification of microvascular sO_2_ for label-free optical imaging oximetry.

In this work, we develop a new data-driven deep spectral learning (DSL) method to enable highly robust and reliable sO_2_ estimation. By training a neural network to directly relate the spectral measurements to the corresponding independent sO_2_labels, DSL bypasses the need for a rigid parametric model, similar to existing deep learning methods for solving optical inverse problems[33–37]. We show that DSL can be trained to be highly robust to multiple sources of variabilities in the experiments, including different devices, imaging protocols, speeds, and other possible longitudinal variations.

A crucial feature of our DSL method is uncertainty quantification. Due to biological variations and tissue heterogeneity, an assessment of the reliability of each sO_2_ measurement is crucial in clinical applications and for guarding against vulnerabilities in making overly confident predictions when imaging rare cases[38]. Existing model-based methods generate a single value of sO_2_ for each spectral measurement, i.e. a point estimate. The accuracy and uncertainty of the point estimate can only be assessed by taking repeated measurements against the ground-truth in a well-controlled experiment. This uncertainty estimation presents aclear limitation for many biomedical applications in which the ground-truth is often inaccessible *in vivo*, and the statistical analysis can only be performed retrospectively on those repeated measurements [13,19,22,32,39]. Instead of assessing the variabilities in the data retrospectively, we develop our DSL model based on an uncertainty learning framework[37] to encapsulate the statistics in the learned model, essentially shifting the burden of repeated measurements in the model-based methods to the training phase of DSL. After the training, the DSL model predicts both sO_2_ and its tandem standard deviation, assessing the uncertainty for each sO_2_ prediction (i.e. a statistical distribution describing all the possible sO_2_ levels of each prediction given the measurements). Most importantly, we show that the DSL predicted statistics closely match those obtained from ensemble calculations. This means that the confidence level calculated from the DSL prediction can be used as a surrogate estimate to the true accuracy of the estimate – making DSL highly reliable.

We demonstrate DSL using two sets of vis-OCT spectral measurements for oximetry on rat retina from [13,19]. Two DSL models are investigated, including a 1D fully connected neural network (FNN) and a 1D convolutional neural network (CNN). Our results show that both DSL models are significantly outperforming the LSF, both in terms of the estimation accuracy and the robustness to experimental variations. We further conduct quantitative statistical analysis based on the uncertainty learning to establish the confidence level of the two proposed models, and further justify the reliability of DSL. Next, imaging oximetry is demonstrated on *en face* sO_2_ maps of rat retina along with the corresponding uncertainty maps, providing a visualization of the DSL predictions. This allows us to assess the accuracy of the prediction based on the underlying physiological conditions.

## 2. Principle of Least Square Fitting

Vis-OCT uses ballistic photon and coherence gating to localize the optical signal within a tissue volume. At the bottom of the vessel wall, light double-passing through the vessel lumen gives rise to the detectable spectral contrast that can be analytically formulated based on the Beer’s law [13]

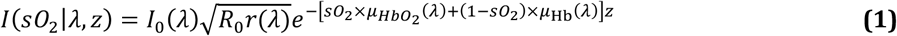

where *I*_*0*_(λ) is the light source’s spectrum; *R*_0_ is the reflectance of the reference arm and assumed to be a constant; r(λ) (dimensionless) is the reflectance at the vessel wall, whose scattering spectrum can be modeled by a power law under the first Born approximation *r*(*λ*)=*Aλ*^−*α*^ with *A* being a dimensionless constant and *α* modeling the decaying scattering spectrum from the vessel wall; The optical attenuation coefficient *µ* (mm−1) combines the absorption (*µ*_*a*_) and scattering coefficients (*µ*_*s*_) of the whole blood, which are both wavelength- and sO_2_-dependent:

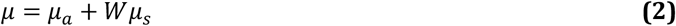

where *W* is a scaling factor for the scattering coefficient and is empirically set to 0.2 [6,21,40]. The subscripts Hb and HbO_2_ denote the contribution from the deoxygenated and oxygenated blood, respectively. *z* denotes the light penetrating depth.

To estimate sO_2_, the traditional approach applies the least-squares procedure that fits the vis-OCT spectral measurement to the analytical model by minimizing the total squares of the error, as illustrated in Fig. 1(a):

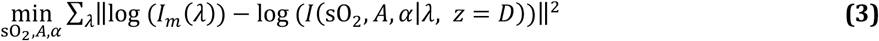

where *I*_*m*_ is typically taken as the vis-OCT spectral measurements extracted from the bottom of the vessel in order to maximize the spectroscopic contrast.

**Fig. 1.**
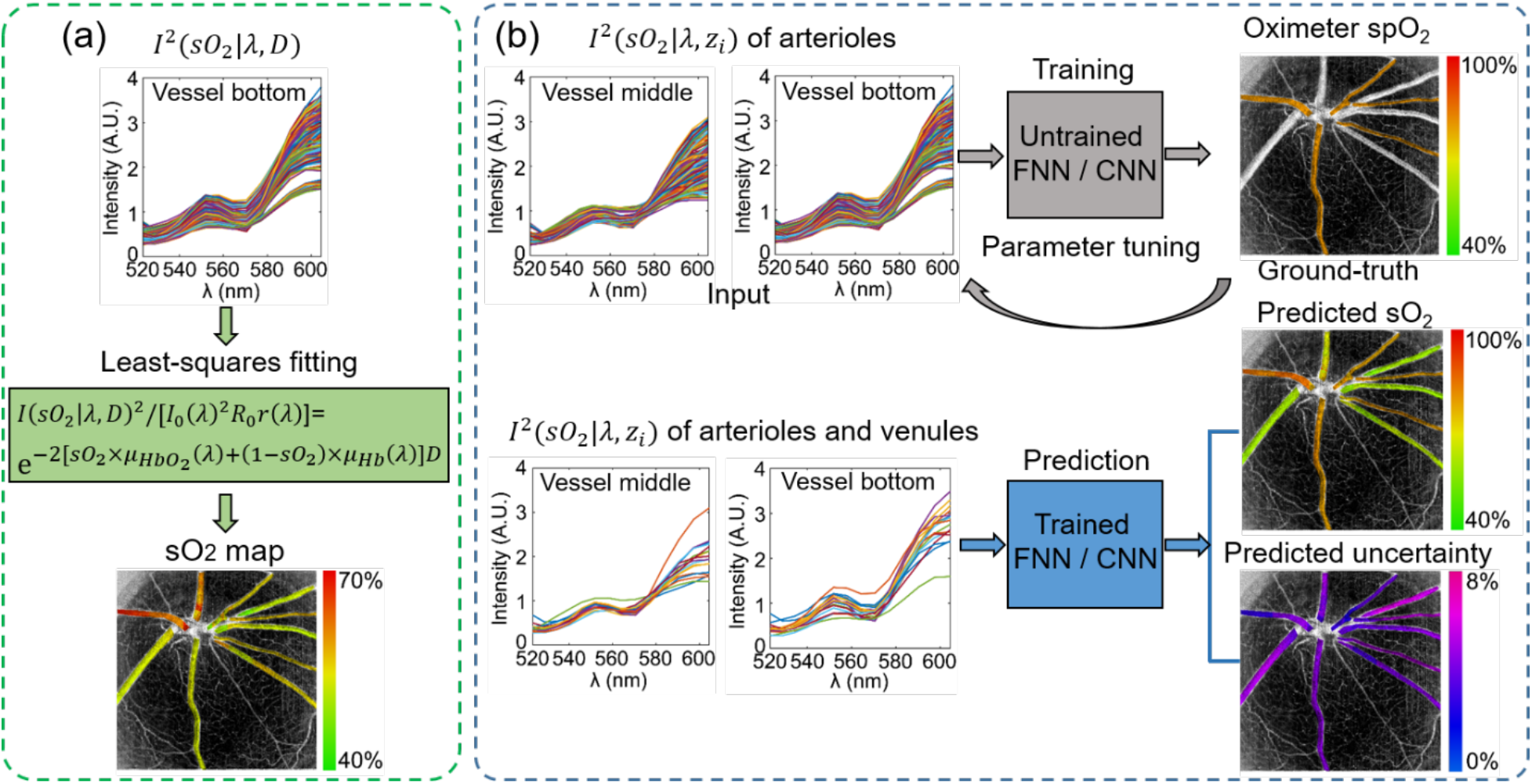
Methods for calculating retinal blood vessel sO_2_ by (a) the traditional LSF, and (b) our neural-network-based DSL with uncertainty quantification. Traditional LSF optimizes a rigid parametric model to best fit the spectral measurements for sO_2_. DSL bypasses any rigid models and trained neural networks with paired arterial spectra and oximeter spO_2_ reading as ground-truth. After training, the neural network models predict both sO_2_ and its uncertainty.

In this analytical model, two free parameters (*A*, *α*) in addition to the unknown sO_2_ level are introduced in order to more accurately capture realistic biophysical interactions. However, in practice these two parameters cannot fully capture all the experimental variabilities. While other models may reduce the free parameters to avoid overfitting [14,17,19,23–25], this approach in general is nonetheless rigid and over-simplified to the actual experiments.

## 3. Principle of Deep Spectral Learning

In DSL, instead of using a rigid analytical model, we train a neural network to link the spectral measurements and the independently measured sO_2_ labels, as illustrated in Fig. 1(b). By doing so, DSL bypasses the need for parametric tuning and model simplification and approximation needed in fitting the analytical models. Furthermore, by removing the restrictions imposed by the analytical model, DSL allows utilizing multiple sets of spectral measurements taken at different depths and enables a more holistic spectral-sO_2_ analysis. Specifically, we demonstrate high-quality predictions using concatenated spectra data from both the bottom and the center of the vessels in vis-OCT. Because the pulse oximeter measures the systemic arterial sO_2_, the same as the retinal arterioles, we use the retinal arterial spectra as the training input paired with the independently measured pulse oximeter sO_2_(spO_2_) as the ground-truth label. After training, the network makes predictions for both arterials and veins.

In addition to sO_2_ prediction, a major important feature of our data-driven DSL method is to quantify the uncertainty for *each* prediction. To do so, we specially design the loss function for training the neural network to properly capture the underlying statistics of the data. The commonly used loss function, such as the mean squared error (MSE), assumes a homogeneous Gaussian distribution of the errors in the predictions. This severely limits its ability to adapt to different types of spectral data variations (e.g. spectral signal outliers, non-uniform noise, and unevenly sampled data) that are inevitably *in homogeneous*. To account for this, we design a customized loss function derived from a *heterogeneous* Gaussian distribution model. Using the training data set (*I*_*i*_, [spO_2_]_*i*_), *i* = 1,2,…, *N*, where *I*_*i*_ and [spO_2_]_*i*_ are the *i*^th^ vis-OCT spectral measurement and the ground-truth pulse oximeter spO_2_, respectively, our loss function *L*_*G*_(*w*) is

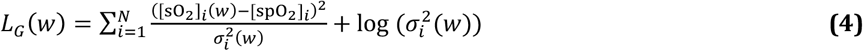

where [sO_2_]_*i*_ and *σ*_*i*_ denote respectively the neural network predicted mean and standard deviation of the underlying Gaussian distribution for the *i*^th^ training data pair; *w* is the learned neural network weights. The main idea of this loss function assumes that the prediction made on each data follows a distinct Gaussian distribution, and the network is trained to predict the underlying mean and standard deviation [41]. Accordingly, the standard deviation σ quantifies the uncertainty for each sO_2_ prediction.

We investigate two neural network models, including an FNN, anda CNN model, whose network architectures are shown in Fig. 2(a) and 2(b), respectively. The detailed descriptions of the data preprocessing and network architecture are included in the method section.

**Fig. 2.**
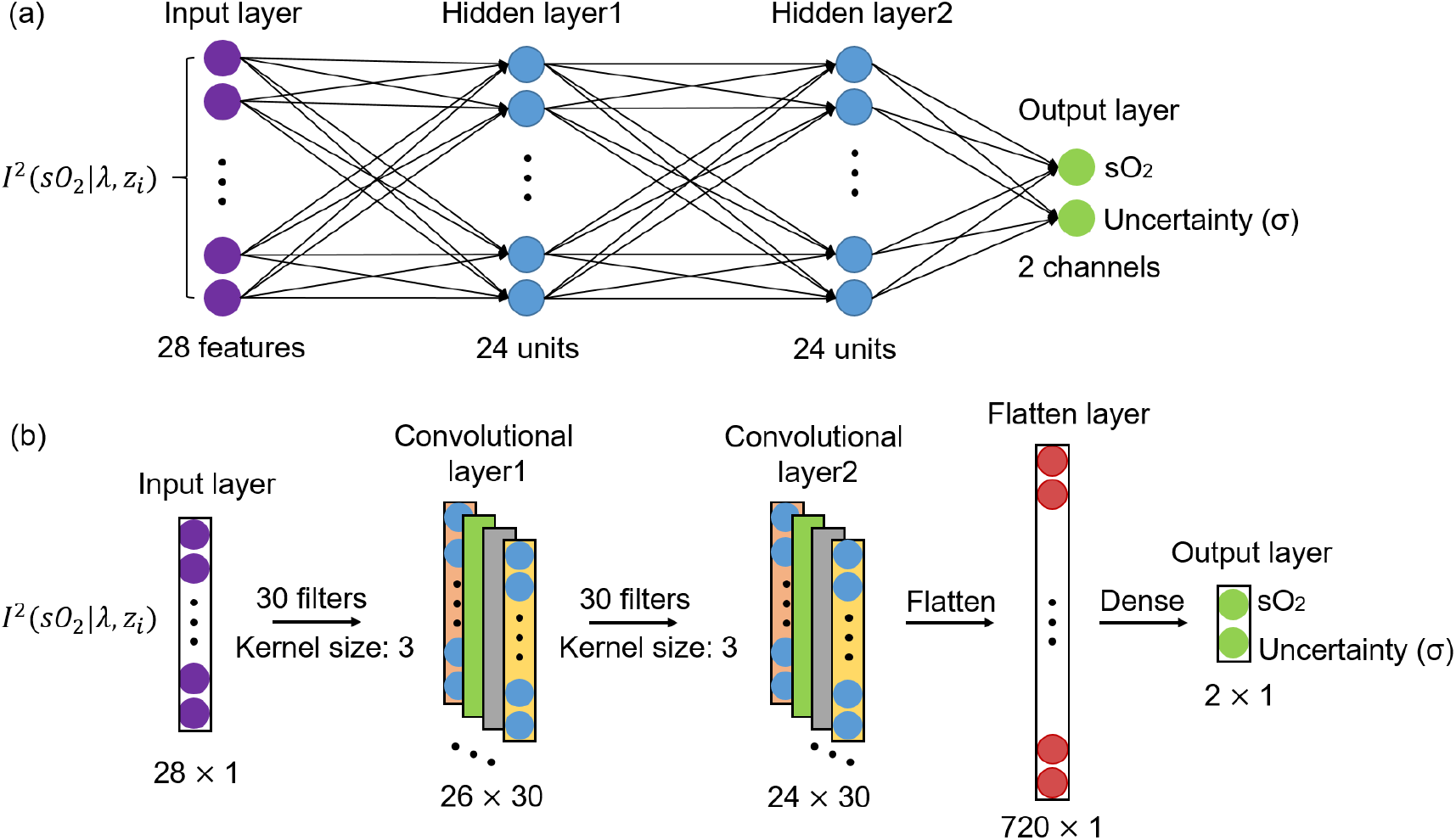
Structures of the FNN model (a) and the CNN model (b) for sO_2_ prediction with uncertainty quantified by the predicted standard deviation.

## 4. Results and Discussion

### A. Data source

To evaluate the effectiveness of the DSL approach, we compiled two datasets from the previous literatures on vis-OCT for retinal oximetry [13,19]. Specifically, the data in Fig. 3(a) are from Ref. [13]and Fig. 3(b) from Ref. [19]. Both data sets used similar experimental protocols that the oxygen content in the ventilation gas were adjusted to induce systemic hypoxia or hyperoxia, and vis-OCT measurements were taken at each ventilation condition. In Ref. [13], a step-wise hypoxia challenge was given to four rats, reducing oxygen content gradually from 21% to 9%. In Ref. [19], eight rats were measured by five ventilation conditions, sequencing from normoxia, hyperoxia (100% O_2_), 5% carbon dioxide (21% O_2_, 74% N_2_, and 5% CO_2_), hypoxia (10% O_2_, 90% N_2_), and finally to normoxia. At each ventilation condition, the systemic arterial spO_2_ reading was taken by a pulse oximeter attached to a rear leg of the rats. The spO_2_ readings are used as the ground-truth label for the major retinal arterioles for neural network training. Depth-dependent backscattering spectra of rat retinal arterioles in vis-OCT were extracted at each ventilation conditions as spectral input to the neural network. The extracted arteriole spectra with the spO_2_ labels were then split into training and testing sets. In Ref. [13], data from three of the four rats were used as the training sets with the remaining one as the testing set. In Ref. [19], data from seven rats were used as the training sets and the remaining one is the testing set. Fig. 3(a) and 3(b) summarize the number of spectra extracted and the corresponding spO_2_ labels in descending order, respectively. Fig. 3(c) shows the histogram of all the training and testing spectra with their corresponding spO_2_ labels.

**Fig. 3.**
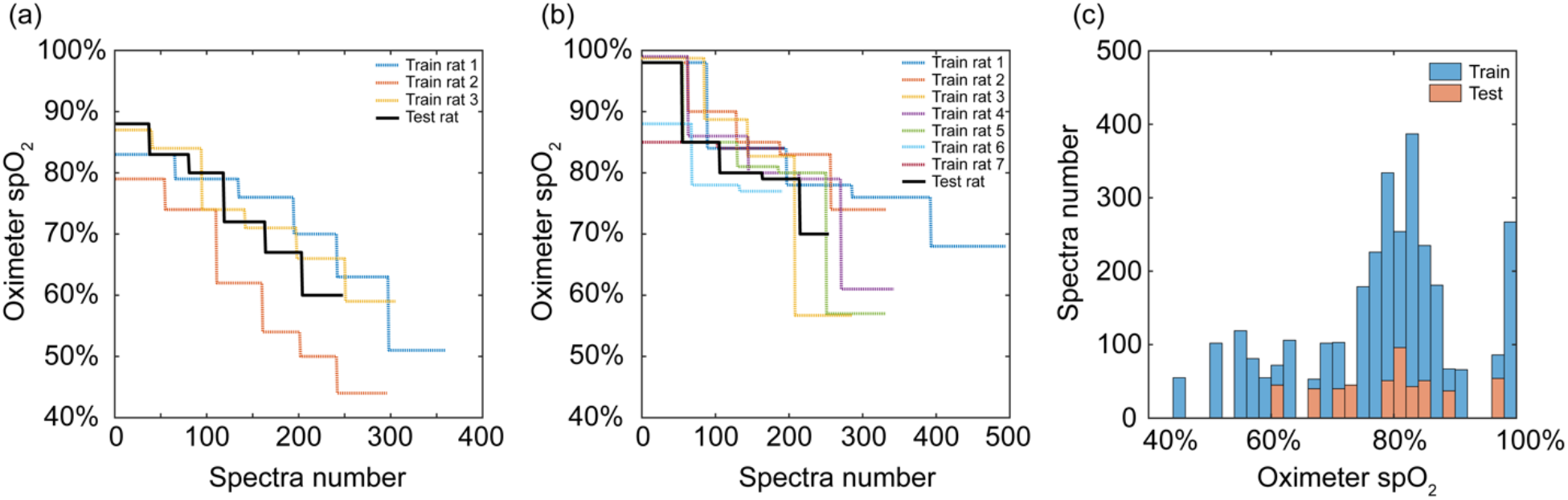
Oximeter spO_2_ readings (ground-truth labels) for training and testing. (a) The readings of rat retinal arterioles from normoxia to hypoxia from Ref. [13]. (b) The readings of rat retinal arterioles from hyperoxia, 5% CO_2_, normoxia to hypoxia from Ref. [19]. (c) The histograms of all the readings for both training and testing sets.

### B. Prediction of arterial sO_2_

Once the networks are trained, the sO_2_ prediction from the testing data by FNN and CNN are plotted in Fig. 4(a) and 4(b), respectively, along with the ground-truth oximeter spO_2_ readings and the LSF estimates for comparison. The first 248 testing data are from Ref. [13] and the rest 254 are from Ref. [19]. Using the data from Ref. [13], the sO_2_ estimation using both DSL models (FNN, CNN) and the LSF agree well with the change in the spO_2_ readings. Much lower variations are observed in the DSL predictions than those from LSF, owing to DSL’s improved robustness to noise and other random signal fluctuations. When the same LSF parameters and inverse calculation procedure for the data from Ref. [13] are applied to the data from Ref. [19], large deviations from the oximeter spO_2_ readings are observed in the estimates. In comparison, the sO_2_ predicted by both DSL models consistently agree with the spO_2_ readings, demonstrating the DSL’s robustness to experimental variations present in these two data sets, such as difference in SNR, imaging protocol and speed. The absolute errors between the sO_2_ predictions and the corresponding spO_2_ from the FNN and CNN are plotted in Fig. 4(c) and 4(d), as compared to those from the LSF. Errors from both DSL models are significantly lower than the LSF model, especially for the testing data from Ref. [19]. To quantitatively compare the three different models, we calculate the mean square errors (MSE) of the FNN and CNN models as 0.3149% and 0.2665%, respectively, both of which are much lower than 3.870% by the LSF. An important feature of our DSL models is its ability to quantify uncertainty via the tandem standard deviation (σ) for each sO_2_ prediction (Fig. 4(e) and 4(f)). Overall, both FNN and CNN predict σ around 5-7%, and the variation of σ justifies the use of heterogeneous Gaussian distribution model in our customized loss function. We also see that the prediction for Ref. [19] has lower variation on σ than Ref. [13], presumably due to more animal numbers and larger training dataset (Fig. 3). Out of 2779 training data, 1930 of them are from Ref. [19].

**Fig. 4.**
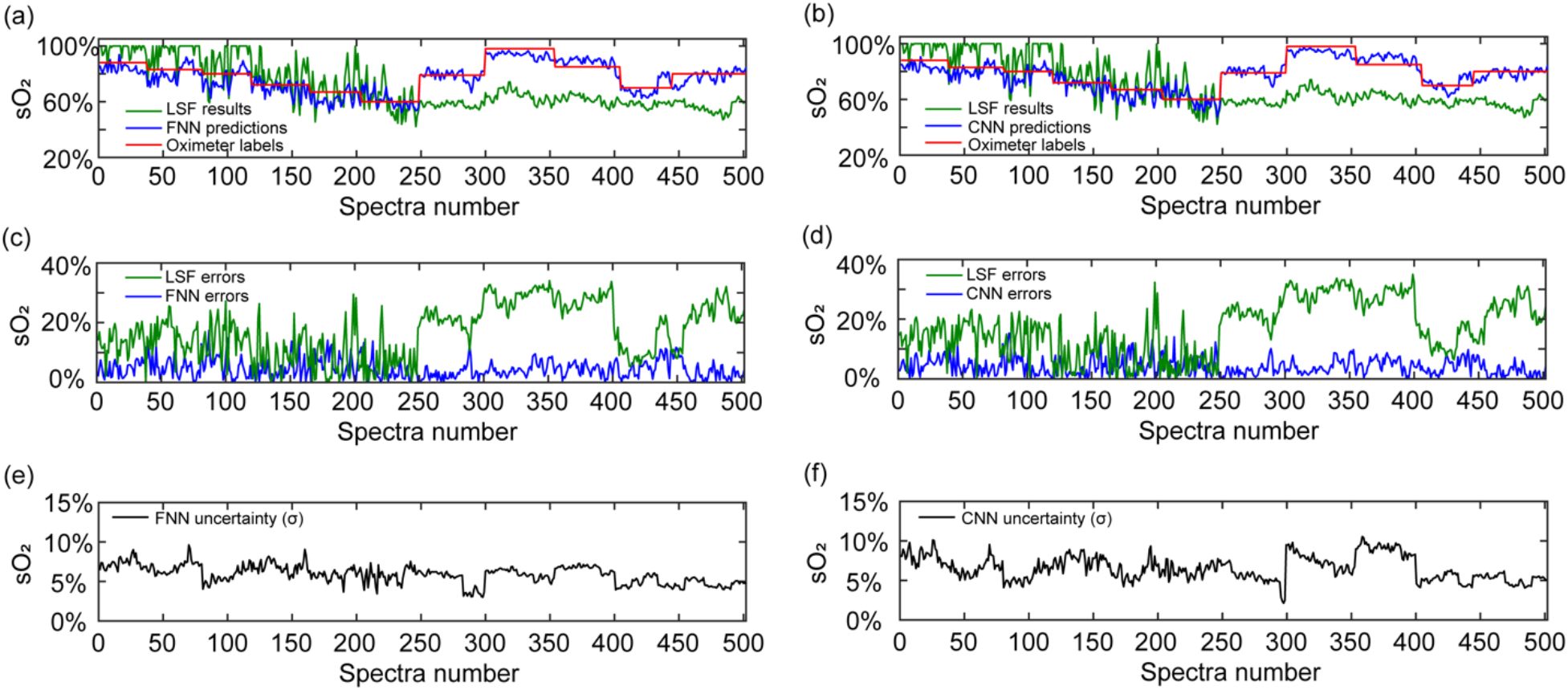
Rat retinal arterioles sO_2_ of the testing data at different ventilation conditions predicted by the FNN, CNN, and LSF models. The predicted sO_2_ by the FNN (a) and CNN (b), compared with the oximeter spO_2_ readings and the LSF calculations. The errors of the predicted sO_2_ by the FNN (c) and the CNN model (d) as compared to the LSF results. The predicted uncertainties of sO_2_ by the FNN (e) and the CNN model (f) measured by the standard deviations (σ).

### C. Evaluation of the quantified uncertainty

Our uncertainty quantification assumes that the predicted sO_2_ follows a heterogeneous Gaussian distribution given different spectral inputs. To validate our uncertainty metrics, we retrospectively calculated the actual probability that the ground-truth (spO_2_) falls within a certain confidence interval of the predicted sO_2_, and summarize the results using the reliability diagram [34,37,42]. To construct a reliability diagram, we gather a sub-set of predictions with aspecific standard deviationσ_0_. We then calculate the probability from this sub-set of data satisfies the criterion of |[sO_2_]i − [spO_2_]i| < *ησ*_0_, where [sO_2_]i and [spO_2_]i are the prediction and the corresponding groundtruth from the ith vis-OCT spectrum:

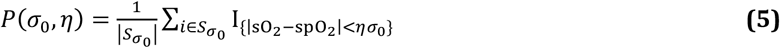

where *η* is a variable that defines the confidence interval, and *S* denotes the sub-set of the prediction with the specified standard deviation. In practice, we relax σ_0_ to σ_0_±1% in order to include sufficient data for ensuring reliable statistical calculations. Intuitively, the probability will approach 1 when *η* increases, *i.e.* a larger error tolerance. At the same time, *η* has a one-to-one correspondence to the theoretical confidence value calculated from the normal distribution. The reliability diagram essentially plots the actual probability against *η*, or theoretical confidence. For an ideal well-calibrated model, the actual probability *P*(*η*_0_, *σ*) should equal to the theoretical confidence - falling on the diagonal line in the graph. When the actual probability *P*(*η*_0_, *σ*) is lower than the theoretical confidence, it indicates that the model is over-confident - the reliability curve is under the diagonal line. When the actual probability *P*(*η*_0_, *σ*) is higher than the theoretical confidence, it indicates that the model is conservative - the reliability curve is over the diagonal line.

The reliability diagrams for both models are shown in Fig. 5. To cover over 90% of the total 502 predictions of the testing data in the reliability diagram, we set σ_0_ = 5% and 7% for FNN model and σ_0_ = 5%, 7% and 9% for the CNN model, respectively. For both models, the sO_2_ predictions with uncertainty falling within the 7%±1% range are slightly conservative, with *P*(*σ*_0_, *η*) higher than the predicted confidence; for the FNN model, the sO_2_ predictions with uncertainty σ0 = 5% are slightly over-confident since *P*(*σ*0, *η*) is lower than the confidence (Fig. 5a); while for the CNN model, results falling within 5%±1% are quite close to the diagonal line (Fig. 5b), indicating a well-calibrated DSL model in this regime. In Ref. [13], the accuracy for sO_2_ calculated by LSF was estimated to be within #x00B0;4% relative error in a well-controlled *in vitro* blood calibration experiment, using blood analyzer readings as the ground-truth. The uncertainty predicted here (∼5% sO_2_) by DSL agrees reasonably well with the *in vitro* calibration result.

Supplemental Fig. 1 illustrates the sub-sets of data when σ_0_ = 5%, 7% and 9% for both models. Table 1 provides a summary of the fitting parameters, slope and constant, and how many of the total 502 testing data were counted for the statistical analysis within each uncertainty range.

**Fig. 5.**
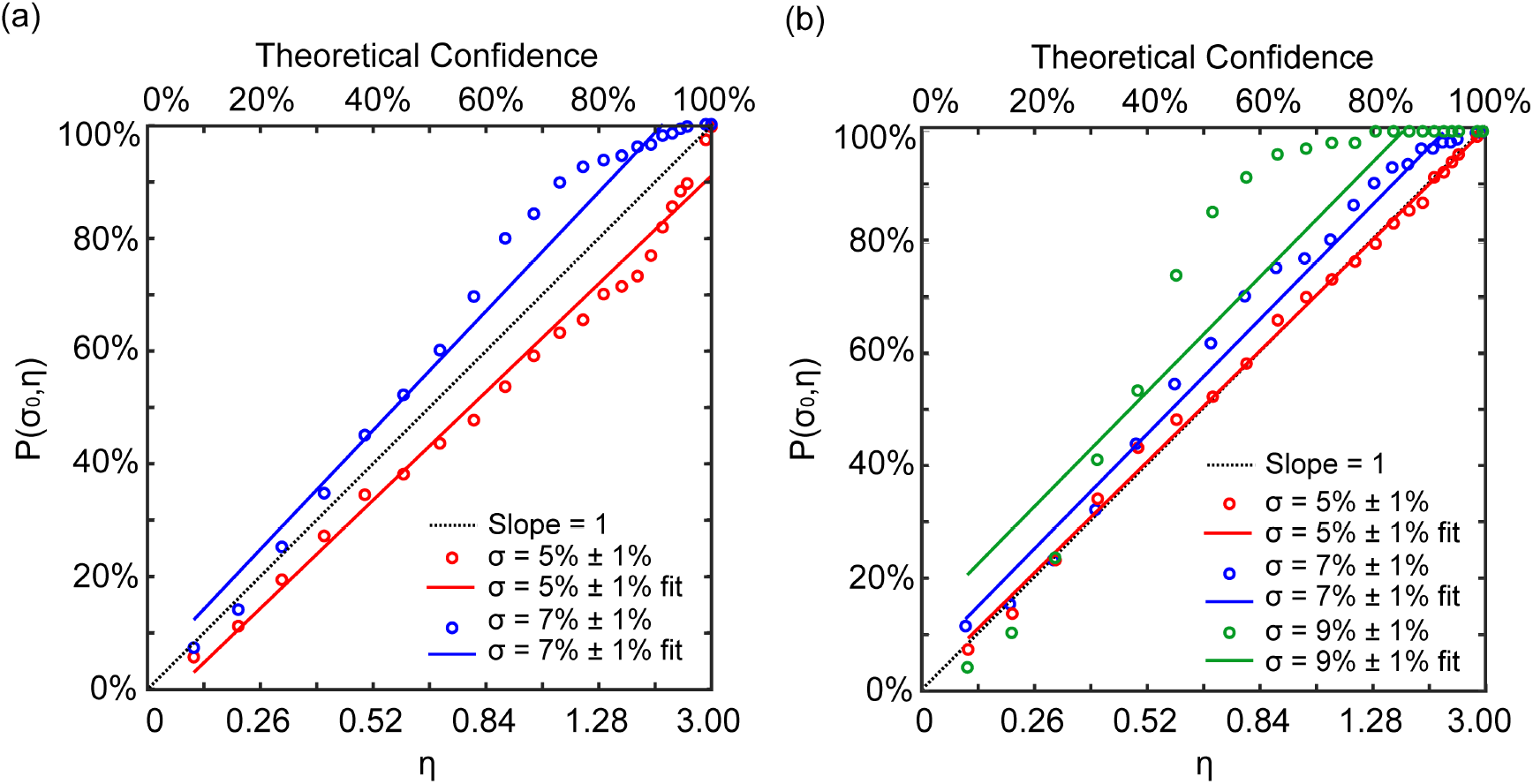
Statistical analysis of the sO_2_ predictions with the quantified uncertainty. (a) The Linear fit of the empirical accuracy to the confidence for the FNN model when the uncertainty measured by the standard deviation (σ_0_) is 5% and 7% sO_2_. (b) The linear fits of the empirical accuracy to the confidence for the CNN model when the standard deviation (σ_0_) is 5%, 7%, and 9%.

**Table 1:** Parameters of linear fit of *P*(*σ*_0_, *η*) to theoretical confidence for sO_2_ predictions with different uncertainties (*σ*) by the FNN and the CNN models.

| Models and uncertainties | Slope | Constant | Datasets |
| --- | --- | --- | --- |
| The FNN model ($\sigma: 5\% \pm 1\%$) | 0.9600 | -0.04886 | 219 |
| The FNN model ($\sigma: 7\% \pm 1\%$) | 1.057 | 0.03658 | 252 |
| The CNN model ($\sigma: 5\% \pm 1\%$) | 0.9878 | 0.01440 | 219 |
| The CNN model ($\sigma: 7\% \pm 1\%$) | 1.0233 | 0.04901 | 178 |
| The CNN model ($\sigma: 9\% \pm 1\%$) | 1.0229 | 0.1250 | 97 |

### D. *En face* sO_2_ maps with uncertainty quantification

After model test and uncertainty analysis, we used the testing data from Ref. [19] and applied the FNN and CNN models for retinal imaging oximetry in comparison to LSF (Fig. 6), at three different ventilation conditions. The oximetry by both FNN and CNN clearly reflect the sO_2_ changes of all vessels from hypoxia, normoxia, to hyperoxia, with the predicted sO_2_ of arterioles matching with the oximeter spO_2_ readings well. The sO_2_ contrast between arterioles and venules are also clearly visualized in hypoxia and normoxia. In comparison, although the sO_2_ by the LSF reflects a similar sO_2_ change form hypoxia to hyperoxia and the arteriovenous sO_2_ contrast, all estimated arterioles values are much lower than the spO_2_ readings due to the experimental variations present in the two experiments. There are also higher variances in the sO_2_ results within each individual blood vessel by LSF and DSL, particularly in Fig. 6(g) and Fig. 6(i). These results clearly indicate the superior robustness and resilience to variations in experimental conditions and within vessels by DSL. Importantly, our DSL models enable direct visualization of the “*en face*” uncertainty maps of the sO_2_ predictions in Fig. 7. The FNN and the CNN models have similar uncertainty estimation on the sO_2_ predictions (Fig. 7(a)–7(f)). At hypoxia, it appears the sO_2_ estimation is inconsistent at the periphery on the right side of the images, as pointed out by the black arrows in Fig. 6(a), and 6(d). At those regions, both DSL model predicted *σ* < 4% (Fig. 7(a) and 7(d)). Combined with the analysis in Fig. 5, it suggests that one should be cautious due to the over-confident predictions.

**Fig. 6.**
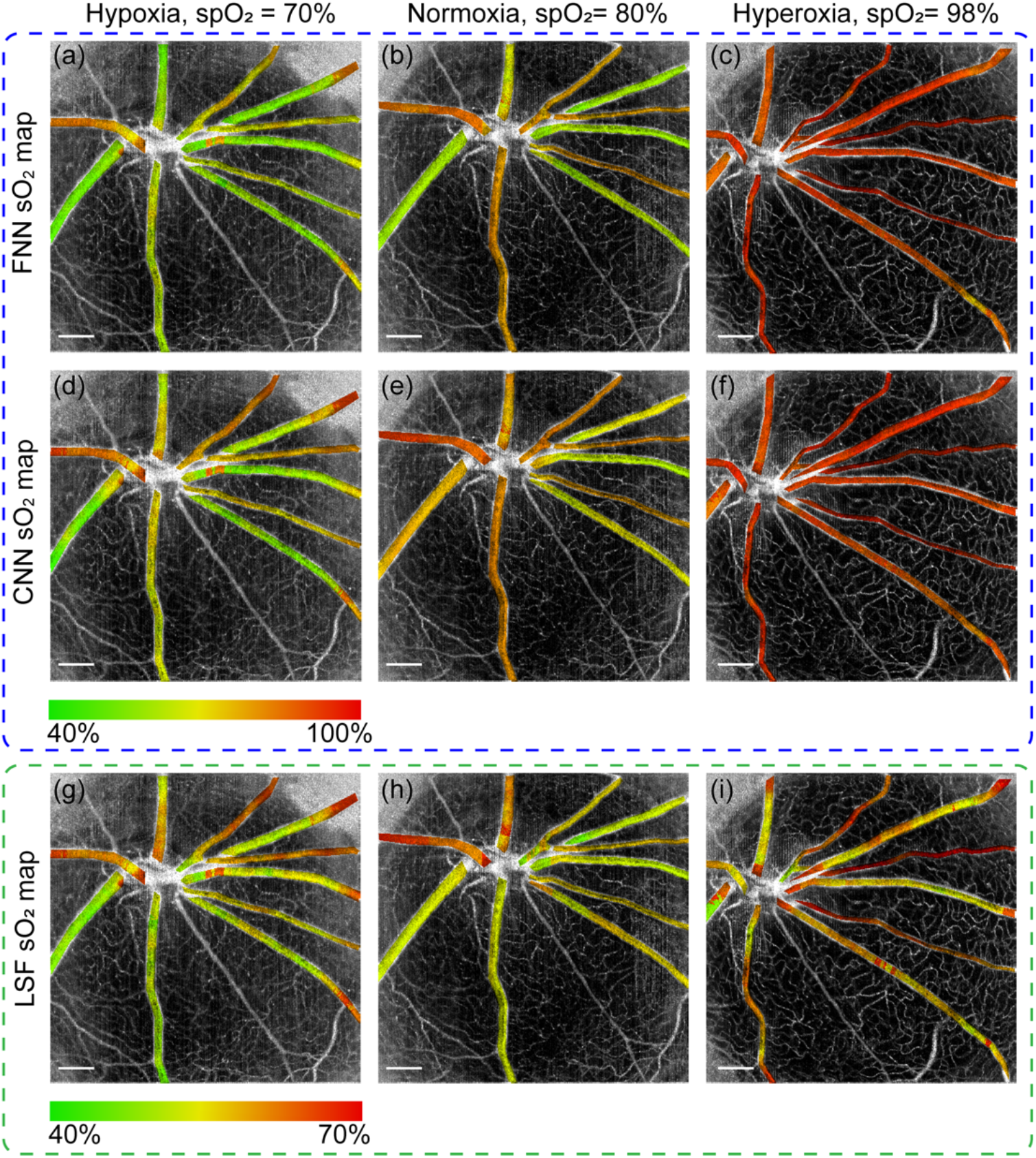
The *en face* sO_2_ maps of the testing data for rat retinal oximetry by FNN (a-c), CNN(d-f) and LSF(g-i), at hypoxia in (a), (d) and (g), normoxia in (b), (e) and (h), and hyperoxia in (c), (f) and (i). The oximeter spO_2_ readings at hypoxia, normoxia, and hyperoxia status are 70%, 80%, and 98%, respectively. Black arrows in (a) and (d) points to regions having inconsistent sO_2_ predictions from other regions in the same vessel. Scale bar: 500 μm.

**Fig. 7.**
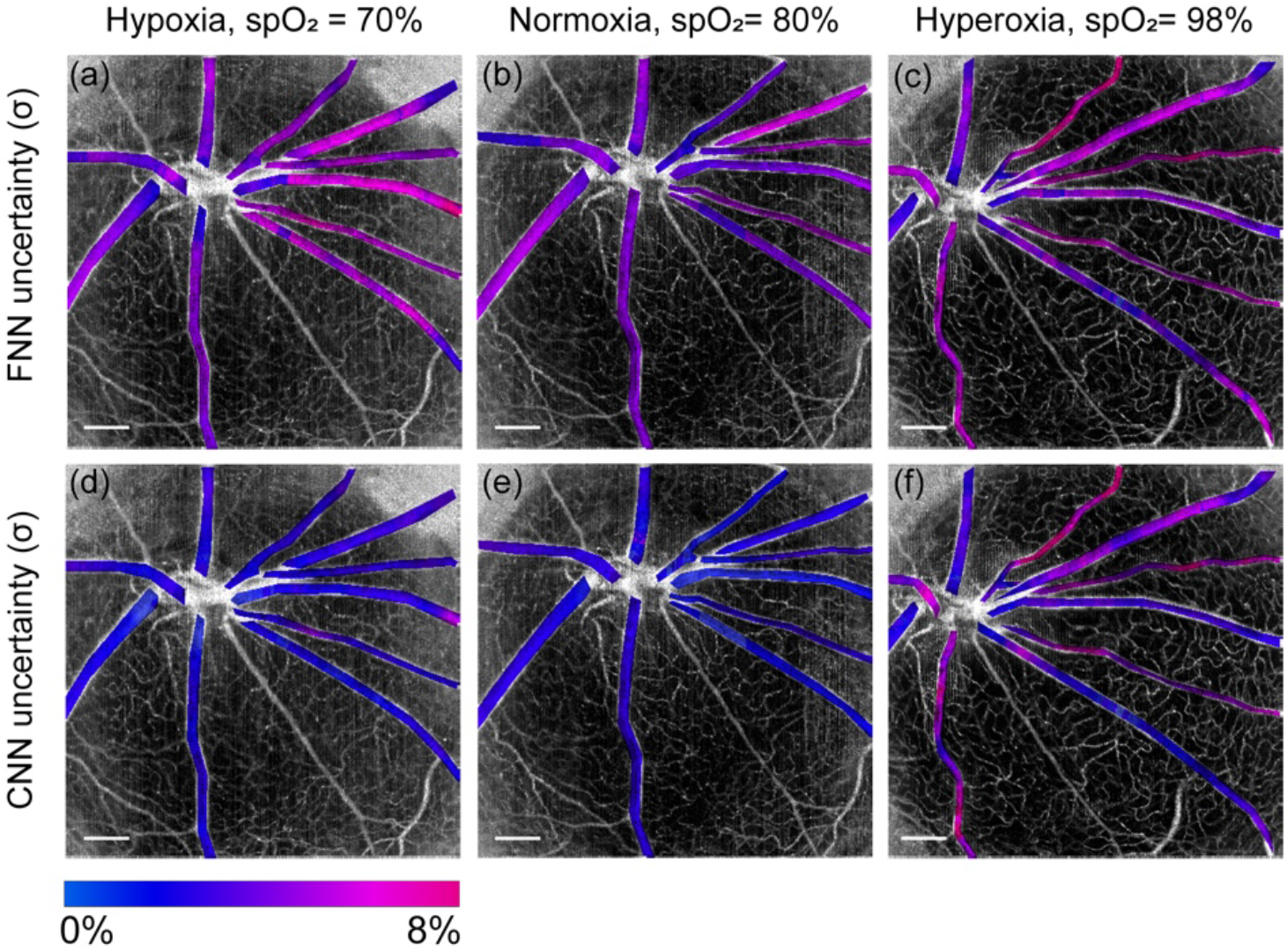
The *en face* uncertainty (σ) maps for sO_2_ predictions corresponding to Fig. 6(a)-6(f). (a-c) Uncertainty maps predicted by the FNN model at three ventilation conditions. (d-f) Uncertainty maps by the CNN model at three ventilation conditions. Black arrows in (a) and (d) points to regions having inconsistent sO_2_ predictions from other regions in the same vessel. Scale bar: 500 μm

## 5. Summary

In this paper, we present a new framework for optical imaging oximetry based on deep spectral learning (DSL). The DSL offers several unique advantages as compared to existing least-square-fitting (LSF) method. First, it bypasses the need for any rigid analytical models and is highly flexible and resilient to experimental variations. We tested DSL on two datasets from two separate vis-OCT experiments, and showed that DSL maintained consistent agreements with the ground-truth spO_2_ despite many differences in these two experiments. In contrast, LSF with the identical parametric setting generated significant biases between the two datasets. Second, without the restriction of any rigid models, DSL allows more flexible and efficient use of the data. Here, we demonstrate the effectiveness of using the spectra from both the middle and bottom of the vessels, since both carry sO_2_-dependent spectral contrast. Most importantly, DSL not only provides the point estimate of sO_2_ but also quantifies the tandem uncertainty of the prediction. Quantifying the statistical uncertainty for each measurement is not possible using the traditional LSF approach, however is valuable in assessing the fidelity of each measurements, in particular in clinical applications. We validate the uncertainty quantification by using the reliability diagram, and for the first time to our knowledge, have constructed uncertainty map of *in vivo* imaging oximetry showing the estimation confidence by DSL. More generally, our DSL framework presents an attractive data-driven approach for other inverse scattering spectral analysis beyond oximetry.

## 6. Methods

### A. Vis OCT experiments

The vis-OCT systems in Ref. [13] and Ref. [19] had the same spectral range from 520 nm to 630 nm, with the same lateral and axial resolutions estimated as 15 *μm* and 1.7 *μm*, respectively. The scanning protocol in Ref. [13] used a raster scan over a 20° square retinal area covering a field of view (FOV) of 2.51mm×2.51mm, with 256×256 pixels in each direction at a 25 kHz A-line rate. The exposure time for the spectrometer camera was 37µs. The entire vis-OCT image stack took 3.3 seconds to acquire. The scanning protocol in Ref. [19] was for optical coherence tomography angiography, scanning a 40 ° square retinal area covering a FOV of 4.37mm × 4.37mm, with 400 pixels in the A-line direction and 512 pixels in the B-scan direction at a 50 kHz A-line rate. The exposure time for the spectrometer camera was 17µs. For the sake of vis-OCT angiography, there were repetitive (5×) unidirectional B-scans of the same cross section, giving a total of 5×512 B-scans for each acquisition. The entire vis-OCT image stack took 25.6 seconds to acquire.

### B. Spectral Extraction and Data Preprocessing

Wavelength-dependent vis-OCT images were first generated by a short-time Fourier transform (STFT) with 14 equally spaced Gaussian spectral windows in the k-space. The wavelength spans from 523.4 – 604.5 nm. The size (FWHM) of the Gaussian window in the k-space is 0.32 μm^−1^, corresponding to a bandwidth of ~17 nm at 585 nm. After STFT, a spectrum can be obtained at each 3D vis-OCT voxel. Next, we performed segmentation to isolate the spectra within retinal arterioles [21,22]. Retinal blood vessels are first segmented from the *en face* projection of the 3D vis-OCT image by a threshold-based algorithm [22]; next all A-lines within the segmented retinal arterioles are shifted in the axial direction in reference to the retinal surface and randomly shuffled. A rolling averaging of 100 shuffled A-lines in Ref. [13]and 250 shuffled A-lines in Ref. [19] was performed with 50 and 125 rolling step size. Because the major vessels are located on top of the retina, signal within the vessels can be averaged in reference to the retinal surface to generate one spectrum. We located the bottom vessel wall[13,18], and averaged signals ±16.63 µm as the vessel bottom spectrum. We then averaged the signal from ~25 µm above the bottom vessel and within a range of ±8.31µm as the spectrum from vessel center. For DSL, these two spectra were concatenated as a whole spectral input. Finally, each individual spectral input was normalized by its mean to ensure similar scaling of all datasets before neural network training.

### C. Network architectures and training

The first FNN model as illustrated in Fig. 2(a) concatenates two spectra from both the bottom and center of the vessels with a size of 1×28 as input. The output predicts both the mean sO_2_ level and the uncertainty (measured by the standard deviation) in two output channels, both of which are single values in a unit of volume percentage of blood being oxygenated. The model has two hidden layers, each having 24 units. We use the Relu activation function in the two inner layers, and the sigmoid activation function in the final layer to normalize the predictions between 0% and 100%.

The second CNN model as illustrated in Fig. 2(b) takes the same input as the FNN model, and also predicts the sO_2_ with uncertainty. The model has two convolutional layers, with each of them having a filter size of 3 and a filter numbers as 48, 36, 24, respectively, and one flatten layer. We use the Relu activation function in the two convolutional layers, and the sigmoid activation function in the final layer.

All the data processing and network training are implemented in Python using TensorFlow/Keras library. Both models were trained with an initial learning rate of 2.5×10^-3^ and we gradually decreased the rate by 1/(1 + αN), where N is the epoch number, and α is a decay rate set to be 0.01. The same total epoch number of 2000 with the same batch size of 50 ensured that the learning curve could reach a plateau. We set the validation split ratio as 0.2 and selected the model with the minimum validation loss as the optimal one for following sO_2_ prediction.

### D. Loss function for uncertainty quantification

In the proposed DSL model denoted by its network weights *w*, the network makes estimation on both the mean and the standard deviation of sO_2_ given the input spectral measurement. Assuming that sO_2_ of all retinal arterioles of a rat at each particular oxygenation status satisfy different Gaussian probability distributions,

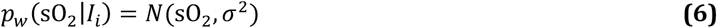

where the mean and the standard deviation of the Gaussian distribution is denoted as sO_2_ and σ, and *N* denotes the normal distribution. The neural network learns a highly-complex nonlinear function. During the training, the network weights, *w*, are estimated by maximizing the joint likelihood over *N* training data pairs:

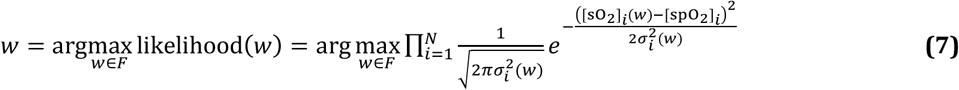

Equivalently, the customized Gaussian loss function *L*_G_(*w*) is minimized when training the DSL models in Eq. 4.

### E. Reconstruction of *en face* Maps of sO_2_ and Uncertainty (σ)

To reconstruct a 2D *en face* map for sO_2_ or uncertainty (σ) we applied the same spatial signal averaging procedure as described in section 7. B., but this averaging was done pixel-wisely within each blood vessel. We first segmented the vessel area manually. For each pixel from the 2D *en face* map within the segmented blood vessel, an arteriole or a venule, its depth-dependent spectra were generated by averaging spectral signals from its 100 (in the first literature) or 250 (in the second literature) nearest neighbors based on Euclidian distance. Then, these spectra would be input into the FNN or CNN model to predict a sO_2_ with uncertainty for this pixel. Next, the above two steps would be iterated pixel-wisely until all pixels within this particular blood vessel had predicted sO_2_ and uncertainty. Finally, the above three steps would be iterated until all arterioles and venules of the rat retina had predicted sO_2_ and uncertainty for displaying their 2D *en face* maps. To generate Fig. 6 and Fig. 7, we applied an algorithm based on the HSV (hue, saturation, value) color model, where the predicted sO_2_ or uncertainty corresponds to image hue, and the angiography signal intensity corresponds to image value and saturation.

## ACKNOWLEDGEMENT

We acknowledge Siyu Chen providing the original data for his vis-OCT experiment. We acknowledge Robert Linsenmeier, and Hao F. Zhang for their supports in both vis-OCT studies.

