## Supplementary material for "Deep spectral learning for label-free optical imaging oximetry with uncertainty quantification": Visualization 1 (from another version of this preprint: version 2, 2019-05-25)

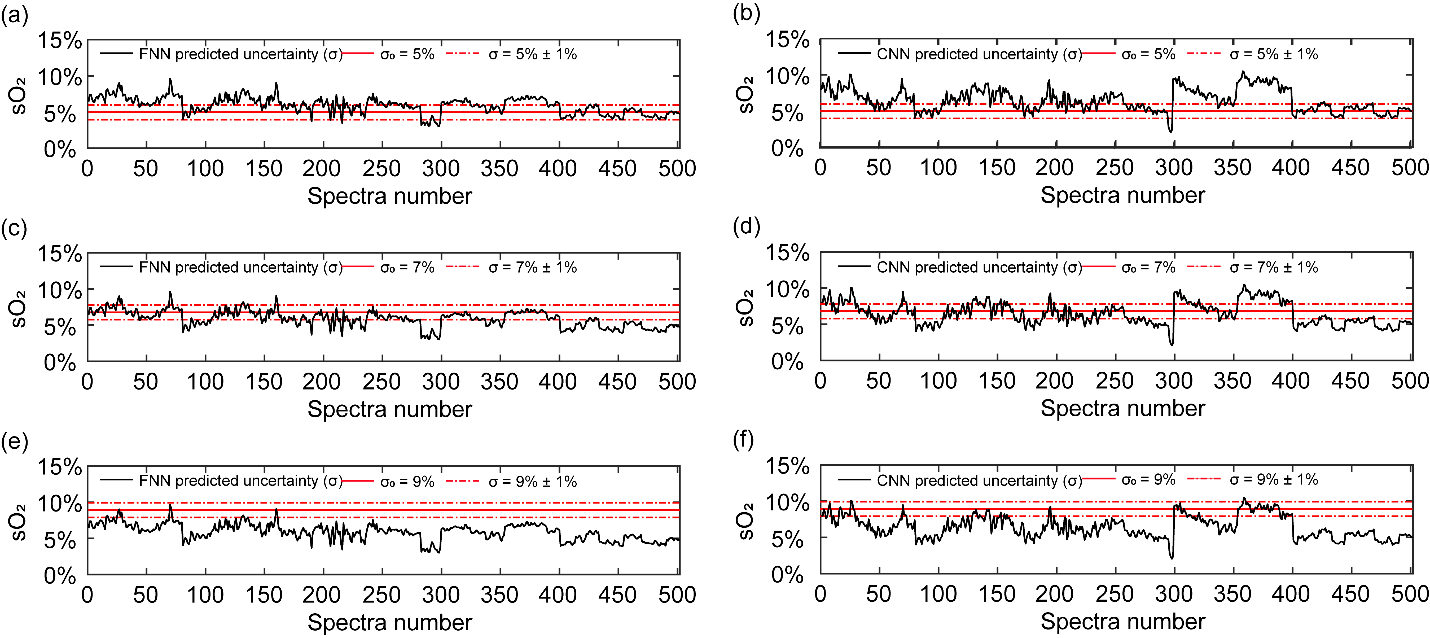

Visualization 1: Illustrations of the sub-set of data when σ_0_ = 5%±1%, 7%±1% and 9%±1%. The FNN model: (a), (c), and (e); the CNN model: (b), (d), and (f).
